# Reward prediction error modulates saccade vigor

**DOI:** 10.1101/555573

**Authors:** Ehsan Sedaghat-Nejad, David J. Herzfeld, Reza Shadmehr

## Abstract

Movements toward rewarding stimuli exhibit greater vigor, i.e., increased velocity and reduced reaction-times. This invigoration may be due to release of dopamine before movement onset. Dopamine release is strongly modulated by reward prediction error (RPE). Here, we generated an RPE event in the milliseconds before movement onset and tested whether there was a causal relationship between RPE and vigor. Human subjects made saccades toward an image. During the execution of their primary saccade, we probabilistically changed the position and content of the image. This led to a secondary saccade following completion of the primary saccade. We focused on properties of this secondary saccade. On some trials, the content of the secondary image was more valuable than the first image, resulting in a +RPE event that preceded the secondary saccade. On other trials, this content was less valuable, resulting in a -RPE event. We found that reaction-time and velocity of the secondary saccade were affected in an orderly fashion by the magnitude and direction of the preceding RPE event: the most vigorous saccades followed the largest +RPE, whereas the least vigorous saccades followed the largest -RPE. Presence of the secondary saccade indicated that the primary saccade had experienced a movement error, inducing trial-to-trial adaptation: the subsequent primary saccade was changed in the direction of the movement error in the previous trial. However, motor learning from error was not affected by the RPE event. Therefore, reward prediction error, and not reward per se, modulated vigor of saccades.

**Author summary:** Does dopamine release before onset of a movement modulate vigor of the ensuing movement? To test this hypothesis, we relied on the fact that RPE is a strong modulator of dopamine. Our innovation was a task in which an RPE event occurred precisely before onset of a movement. We probabilistically produced a combination of large or small, negative or positive RPE events before onset of a saccade, and observed that the vigor of the saccade that followed carried a robust signature of the preceding RPE event: high vigor saccades followed +RPE events, while low vigor saccades followed -RPE events. This suggests that control of vigor is partly through release of dopamine in the moments before onset of the movement.

## Introduction

We tend to move faster toward stimuli that we associate with greater value. For example, when the expected reward following successful completion of a movement is large, saccades [1–3] and reaching movements [4] both exhibit a shorter reaction-time and a higher peak velocity. That is, the reward magnitude that the brain expects to acquire following completion of a movement appears to invigorate the movement. This link between expected reward and movement vigor may be partly due to the function of the basal ganglia [5] and release of dopamine [6], raising the possibility that before every movement, the dopamine that is released in response to the stimulus partly controls the vigor of the ensuing movement.

Dopamine release appears to follow a simple rule. When the acquired reward is unexpectedly large, the neurons fire a burst, but if the same reward is expected, the neurons no longer respond [7,8]. That is, dopamine neurons encode the difference between the predicted stimulus value and the actually acquired value, termed reward prediction error (RPE). This transient encoding of RPE provides an interesting prediction: if dopamine release in the milliseconds before movement onset contributes to control of vigor, then movements that follow a positive RPE (+RPE) should exhibit high vigor, and those that follow a -RPE should exhibit low vigor. That is, vigor modulation should dependent on reward prediction error, not reward itself.

Unfortunately, the hypothesis that RPE (and not reward per se) drives vigor has been difficult to test because in a typical experiment, the RPE event occurs after a movement has been completed and the reward acquired, not before the onset of a movement. Here, we designed a task that overcame this limitation.

In our experiment, we relied on the idea that viewing of images carries some of the hallmarks of reward: when given the option of choosing from various image categories, people prefer face images, and are willing to spend a greater amount of effort in exchange for gaze at those images [9,10]. Furthermore, viewing of face images activates the brain’s reward system [11]. In our experiment we used images as a proxy for reward, and then probabilistically controlled the image content to induce RPE events. We asked whether induction of an RPE event in the milliseconds before a movement influenced vigor of that movement.

Subjects made saccades to view an image, and upon initiation of the saccade, we randomly altered the position and content of that image. The position change forced the subjects to follow their initial saccade with a secondary saccade. Our concern was vigor of this secondary saccade, which usually took place at a latency of less than 150ms with respect to completion of the primary saccade. On some trials, value of image A (primary image) was higher than image B (secondary image), while in other trials value of B was higher than A. As a result, in some trials subjects expected to view a low valued image, but upon completion of their primary saccade, were presented with the opportunity to gaze at a high valued image. This resulted in conditions in which during the milliseconds before the onset of the secondary saccade (as A was replaced by B), there was a +RPE (B>A) or a -RPE (B<A) event. We asked whether the RPE event altered vigor of the ensuing saccade, and found that the secondary saccade carried a robust signature of the preceding RPE event.

## Results

In order to produce an RPE event, we asked subjects to make a saccade toward an image, and then as their movement started, on random trials we replaced that image with one that had a lower or higher value. Notably, we also moved the primary image to a new location, thereby forcing the subjects to follow their primary saccade with a secondary saccade. The replacement of the primary image with a secondary one produced an RPE event that preceded the secondary saccade. We quantified the effects of the RPE event on the characteristics of the secondary saccade.

To produce a +RPE event, the trial began with a noise image (Fig. 1A, left column). As subjects initiated their primary saccade, we probabilistically erased the noise image and replaced it with a face image at a new location (NF trials, first column Fig. 1A). Similarly, in order to produce a -RPE event, the trial began with presentation of a face image, which following saccade onset, was probabilistically replaced with a noise image (FN trials, third column Fig. 1A). The control condition for the +RPE event was a trial in which both the primary and secondary images were faces (FF control, Fig. 1A). The control condition for the -RPE event was a trial in which both the primary and secondary images were noise (NN control, Fig. 1A). Therefore, the secondary saccade in both the control and RPE trials was made toward the same image. However, in the RPE trial, the secondary image was different than the preceding primary image, resulting in what we conjecture was an RPE event. Based on probability of the events, we estimated that the trials produced four magnitudes of RPE (see Methods): highly positive (NF trials), slightly positive (FF trials), slightly negative (NN trials), and highly negative (FN trials).

**Figure 1.**
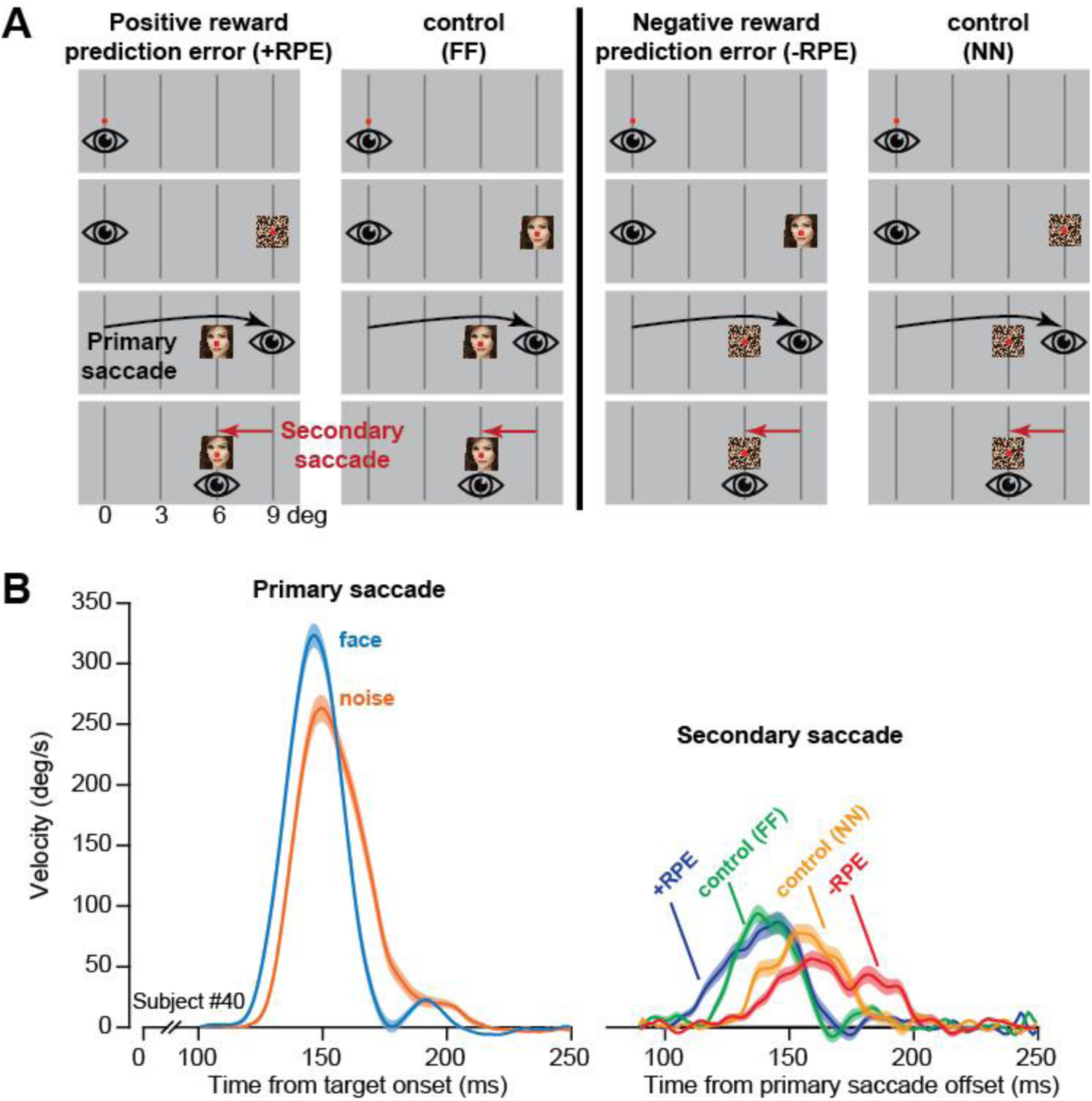
Experiment design and data from a representative subject. **A**. Each trial began with a fixation dot near the center. Following a random fixation interval, we presented a primary image at 9° to the right along the horizontal axis. As the primary saccade took place, we erased the primary image and replaced it with a secondary image. In +RPE trials, a noise primary image was replaced with a face secondary image. A face-face trial served as control for the +RPE trial. In -RPE trials, a face primary image was replaced with a noise secondary image. A noise-noise trial served as control for the -RPE trial. **B**. Saccade velocity for a representative subject. Primary saccade exhibited a shorter reaction-time and a higher velocity in response to a face image. Data for the secondary saccade are plotted with respect to termination of the primary saccade in the same trial. The secondary saccades exhibited shortest reaction-times in +RPE trials, and longest reaction-times in -RPE trials. Error bars are SEM.

### Effects of reward prediction error on vigor

Data from a representative subject are shown in Fig. 1B. As expected, the primary saccade had a shorter reaction-time and higher peak velocity when made toward a face image. During the primary saccade, on some trials the face image was changed to noise (-RPE event, FN trial). Similarly, on some trials the noise image was changed to face (+RPE event, NF trial). Because of the change in image location, at 100-130ms following completion of the primary saccade the subject generated a secondary saccade. We measured the reaction-time of the secondary saccade as latency with respect to end of the primary saccade. The reaction-time and peak velocity of the secondary saccade were affected by not just the image at the destination (i.e., the secondary image), but more importantly, by the sign of the RPE event. Reaction-time of the secondary saccade appeared shortest following the +RPE event, and longest following the -RPE event. Indeed, the properties of the secondary saccade appeared to follow a striking pattern: shortest reaction-time and highest velocity for the most positive RPE event (NF), longest reaction-time and lowest velocity for the most negative RPE event (NF), and in between for the mildly positive (FF) and mildly negative (NN) RPE events.

These results were repeated in our population of subjects. The opportunity to view a face image strongly affected the vigor of the primary saccade (Fig. 2). The probability density of reaction-times shifted earlier (Fig. 2A), from a mean of 150.2 to 140.95 ms, resulting in a within-subject reduction of 9.30±0.63 ms (mean±SEM) (within-subject change, F(1,54)=217, p<10^−4^). The face image also induced an increase in the velocity of the primary saccade (Fig. 2B), particularly in the second half of that movement. As a result, saccade peak velocity increased by 2.63±0.76 °/s (within-subject change, F(1,54)=12.1, p=0.001). Maximum change occurred 10 ms after peak velocity with 6.01±0.84 °/s difference between two categories (within-subject change, F(1,54)=51.511, p<10^−4^). We tried to minimize the changes in saccade amplitude that may arise from changes in image content by presenting a green dot at the center of every image. We observed a very small effect of image type on saccade amplitude: the primary saccades were 8.19±0.01 deg toward face images, and 8.10±0.01 deg toward noise images, a within-subject change of 0.09±0.015 deg (F(1,54)=32, p<10^−4^). However, this change was less than the resolution of our measuring instrument. Overall, the opportunity to view a face image produced a significant reduction in the total time it took for the eyes to arrive at the location of the primary target (Fig. 2C, within-subject change, F(1,54)=199, p<10^−4^).

**Figure 2.**
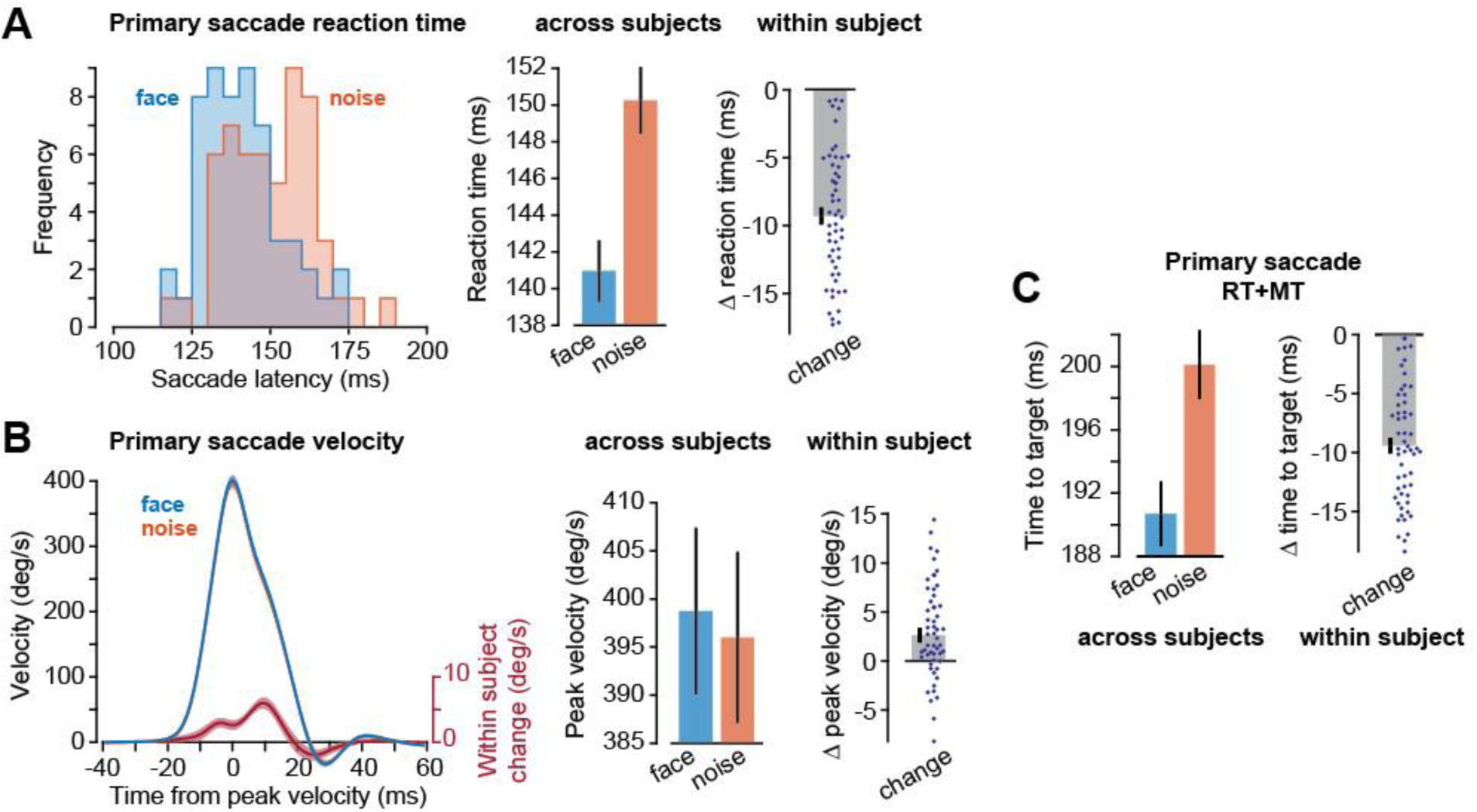
Primary saccades exhibited shorter reaction-time and higher velocity in response to face images. **A**. Distribution of mean reaction-times and within-subject change in the mean reaction-time (face minus noise). Each dot is a single subject. **B**. Saccade velocity traces, and within-subject change in velocity (face minus noise). **C**. Total time to target (reaction-time plus movement duration) and within-subject change. Error bars are SEM.

As the primary saccade started, we displaced the primary image to a new location, encouraging the subjects to produce a secondary saccade. On some trials the content of the secondary image was different from the primary image, potentially causing an RPE event. We found that the reaction-times for the secondary saccade (Fig. 3A and 3B) were shortest in the NF trials (+RPE event), and longest in the FN trials (-RPE event). Indeed, there was an orderly increase in the reaction-times in the precise pattern predicted by the RPE events (Fig. 3B, RM ANOVA F(3,52)=34.7, p<10^−4^). On average, the +RPE event reduced the reaction-time of the secondary saccade by 19.15±1.95 ms with respect to the -RPE event (148.6±3.1 ms in +RPE trial as compared to 167.75±3.3 ms in -RPE trial). Similarly, peak velocity of the secondary saccade was affected by the RPE events (Fig. 3D, RM ANOVA F(3,52)=14.65, p<10^−4^). Amplitude of the secondary saccade on average varied by less than 0.09 deg across the range of the various conditions: +RPE 3.02±0.04 deg, FF 3.08±0.04 deg, NN 3.04±0.05 deg, and FN 3.0±0.05 deg.

**Figure 3.**
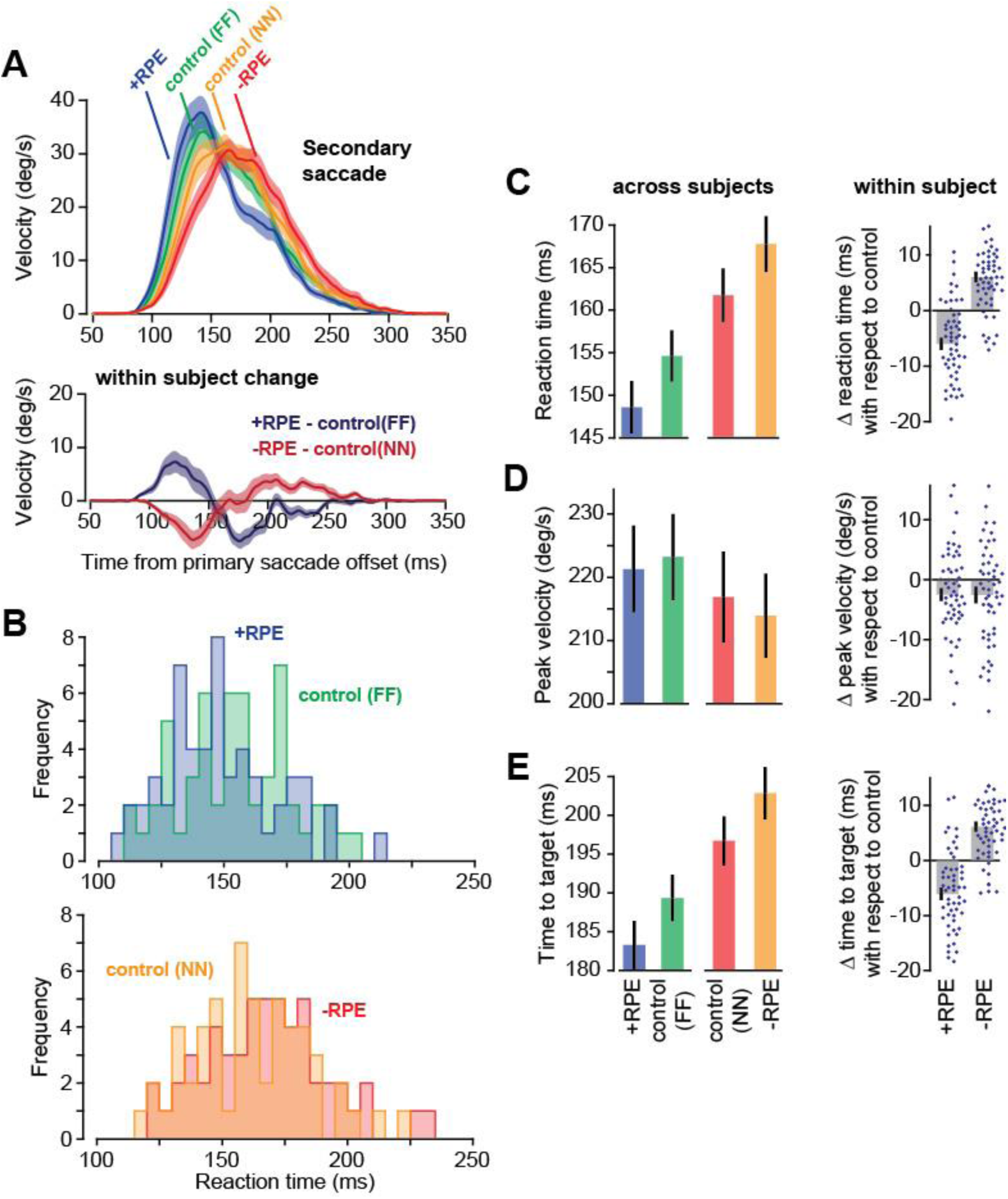
Secondary saccades were influenced by the preceding RPE event. **A**. Saccade velocity and within-subject change in velocity. **B.** Distribution of mean reaction-times across subjects. Reaction-times are measured as the latency with respect to offset of the preceding primary saccade. **C**. Mean reaction-times across subjects and the within-subject change in reaction-times. The bars are +RPE condition with respect to control(FF), and -RPE condition with respect to control(NN). Dots are individual subjects. **D**. Peak velocity and within-subject change in peak velocity. **E**. Total time to target, measured from completion of the primary saccade to conclusion of the secondary saccade, i.e., the reaction-time plus movement duration of the secondary saccade. Error bars are SEM.

Overall, the RPE events significantly affected the total time it took for the eyes to respond and acquire the secondary target (Fig. 3E): the time to target, measured from completion of the primary saccade, was smallest following the +RPE event (NF trials), and largest following the -RPE event (FN trials, RM ANOVA F(3,52)=34.88, p<10^−4^). That is, the magnitude of the RPE event that preceded a saccade affected the vigor of that saccade.

### Effect of reward prediction error on learning

Presence of a secondary (or corrective) saccade indicates presence of a motor error: at the end of the primary saccade, the target was not on the fovea. The resulting movement error should induce plasticity in the cerebellum [12], affecting the subsequent primary saccade. We wondered whether presence of the RPE event modulated learning from the motor error. In particular, would a +RPE event enhance learning?

In our experiment we displaced the primary image along four directions: positive and negative along the horizontal axis (H+ and H-), and positive and negative along the vertical axis (V+ and V-). In all cases the magnitude of the displacement was 3°. Each displacement resulted in a motor error, which in principle may have induced trial-to-trial learning. We measured this learning by comparing the primary saccade that took place before the error to the subsequent primary saccade made to the same visual stimulus type. For example, suppose that on trial *n* the primary saccade was toward face, the subject experienced an H+ error on that trial, and that on trial *n*+1 the primary saccade as again toward face. For all such consecutive pairs of trials, we measured the change in the primary saccade made in trial *n*+1 with respect to trial *n*. This trial-to-trial change in the primary saccade is plotted in Fig. 4A. We found that across all error types, the largest change in the velocity profile was around 15ms after the saccade peak velocity (Fig. 4A). Following an H+ error, the tail of velocity trace (15 ms after peak velocity) increased by 6.62±1.02 °/s (two-sided t-test, t(32)=6.485, p<10^−4^) along the horizontal direction. The trial-to-trial change in saccade amplitude showed 0.17±0.018 deg (two-sided t-test, t(32)= 9.098, p<10^−4^) increase following an H+ error. Similarly, following an H-error, the subsequent primary saccade exhibited a 4.96±1.09 °/s (two-sided t-test, t(32)=4.550, p<10^−4^) decrease in the tail of velocity trace along the horizontal direction and 0.17±0.016 deg (two-sided t-test, t(32)= 10.747, p<10^−4^) reduction in amplitude. Learning was also present following V+ and V-errors. Following a V+ error, there was 1.28±0.42 °/s (two-sided t-test, t(21)=3.038, p=0.006) increase in velocity tail and 0.027±0.012 deg (two-sided t-test, t(21)= 2.348, p=0.029) increase in amplitude of vertical component of primary saccade in the subsequent trial. Following a V-error, there was 1.41±0.42 °/s (two-sided t-test, t(21)= 3.368, p=0.003) decrease in velocity tail and 0.033±0.0086 deg (two-sided t-test, t(21)= 3.809, p=0.001) reduction in vertical amplitude. These results demonstrated that experience of the motor error induced learning, resulting in an error-dependent change in the subsequent primary saccade.

**Figure 4.**
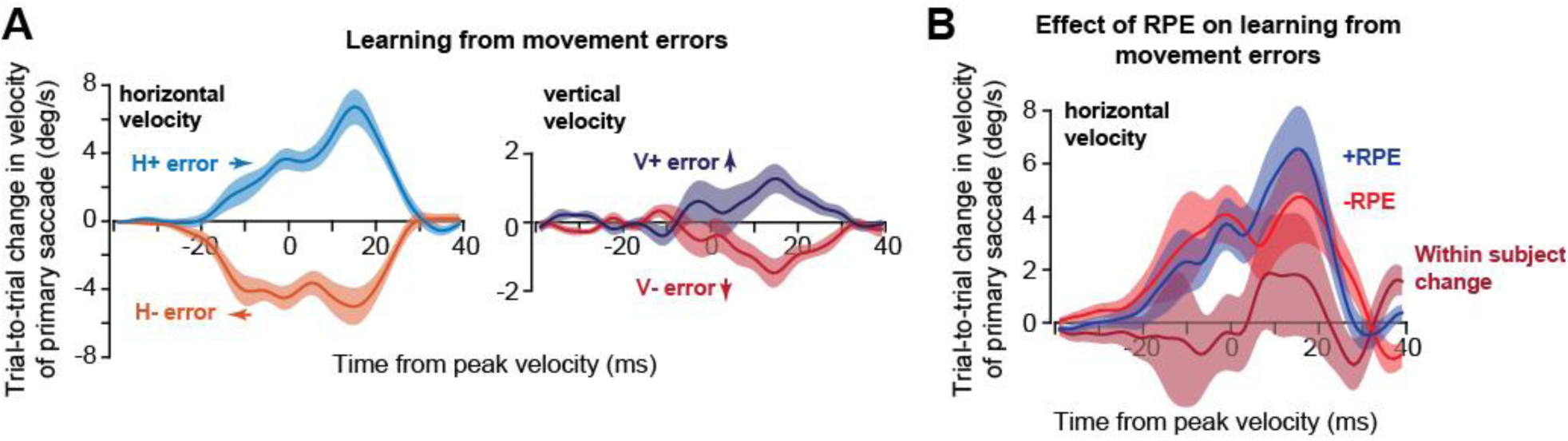
Learning from movement errors and the effect of RPE on learning. **A**. Change in the velocity of the primary saccade from the trial in which the movement error was experienced to the next trial (the primary image type in both saccades were the same). Saccades are grouped based on the movement error experienced at the conclusion of the first primary saccade. Movement error is defined as the position of the secondary image with respect to the primary image. That is, H+ errors imply that the secondary image was further to the right along the horizontal axis than the primary image. The motor error along each axis produced learning along that axis. **B**. Trial-to-trial changes for horizontal motor errors only, grouped based on the RPE event that followed the first primary saccade. Data for H-was flipped and averaged with H+ trials. Within-subject change in learning was marginally larger following a +RPE event (as compared to a -RPE event), but this effect was not significant.

In some trials, a given movement error occurred in the context of a +RPE event, whereas in other trials the same movement error occurred in the context of a -RPE event. To measure the effect of the RPE events on learning, we focused on the horizontal error trials, as the vertical error trials produced substantially less learning. The trial-to-trial change in the primary saccade in the +RPE and -RPE trials are plotted in Fig. 4B. To make this figure, we considered the H+ and H-pairs of trials together (with the positive axis now reflecting trial-to-trial change in velocity along direction of the movement error). We found that learning following a +RPE event (6.46±1.59 °/s change in velocity tail, t(32)= 4.067, p<10^−3^) was marginally stronger than following a -RPE event (4.64±1.73 °/s, t(32)= 2.677, p=0.012), but the effect did not reach statistical significance (1.83±2.32 °/s within-subject change, F(1,32)=0.620, p=0.23).

## Discussion

It is possible that dopamine release in the milliseconds before onset of a movement serves to invigorate the ensuing movement. Here, we attempted to indirectly test this idea by probabilistically changing the potential reward for completing a movement, thereby producing a combination of large or small, negative or positive RPE events. Because dopamine release is affected by the magnitude and direction of the RPE event [7,8], we conjectured that vigor of the movement that followed the RPE event should exhibit a distinct pattern: highest vigor should follow a large +RPE events, whereas lowest vigor should follow a large -RPE event.

Our innovation was a behavioral paradigm in which the RPE events occurred just before the onset of a movement that was required for acquisition of reward. Subjects were presented with the opportunity to view face or noise images. Once they initiated their primary saccade, we probabilistically changed the location and content of the image, resulting in a range of RPEs: highly negative (face change to noise), slightly negative (noise not changed), slightly positive (face not changed), and highly positive (noise changed to face). Following completion of the primary saccade, the subjects experienced the probabilistic change in image position and content, and therefore produced a secondary saccade, which had a latency of around 150ms with respect to termination of the primary saccade. We observed that reaction-times were shortest following the large +RPE event, and longest following the large -RPE event. Saccade peak velocity approximately followed a similar pattern. That is, the magnitude and direction of the RPE event modulated vigor of the ensuing saccade.

Because saccade vigor is dependent on activity of neurons in the superior colliculus [13], and these neurons are influenced by luminance and other low-level properties of the visual stimulus [14], the differences in vigor may have arisen not from presence of an RPE event, but rather because of other variables associated with differences in properties of the secondary image. Therefore, we included control trials in which the saccade of interest was made to the same image as in RPE trials, but without the benefit of an RPE event. We found that in +RPE trials, saccades had shorter reaction-times as compared to image-matched control trials. Similarly, in -RPE trials, saccades had longer reaction-times as compared to image-matched control trials. Furthermore, because the primary and secondary images were always presented at different locations with respect to the fovea, and often in opposite directions, we would expect little or no overlap between regions of collicular activity associated with the primary and secondary saccades. This dissociation between magnitude of primary and secondary saccades reduces the possibility of an interaction between the saccades at the level of colliculus. As a result, it seems likely to us that the vigor differences in the secondary saccades were not solely due to differences in the intrinsic response of the collicular neurons to information that they received from the retina. Rather, the link between RPE events in our experiments and modulation of vigor may have been because of reward-dependent regions that project to the colliculus, such as the basal ganglia and the frontal eye field.

Saccades toward a rewarding stimulus exhibit greater vigor partly because the opportunity for reward reduces the inhibition that the colliculus receives from the substantia nigra reticulata (SNr) [15], and increases the excitation that the colliculus receives from the frontal eye field (FEF) [16,17]. It seems likely that dopamine plays an important role in controlling this drive. Just before onset of a spontaneous movement, there is diversity of responses among dopamine neurons: some show a transient increase, while others show a transient decrease [6]. Notably, for the dopamine cells that increase their activity, the amount of increase is positively corrected with the acceleration of the upcoming movement [6]. In monkeys, dopamine neurons respond to presentation of a saccade target within 100ms, and dissociate between reward and non-rewarding stimuli within 150ms [18]. In our data, saccades that were affected by the RPE event were generated at extremely short reaction-times: around 150ms. Therefore, in principle, the time-range of dopamine response to reward is roughly within the window of the vigor modulation we observed in saccades. Because RPE events have a robust effect on dopamine release, it is possible that the RPE event driven change in saccade vigor is linked through dopamine dependent drive to the basal ganglia and FEF, affecting saccade-related discharge in the superior colliculus.

### Modulation of learning from movement error

In earlier experiments it has been shown that if the primary saccade ends but the target is not on the fovea, the result is a movement error that modulates the probability of complex spikes in the Purkinje cells of the cerebellum [19–21]. The movement error modulation of complex spikes in turn produces trial-to-trial plasticity in the Purkinje cells [12], altering the primary saccade on the subsequent trial. The movement error appears to be transmitted to the cerebellum via a pathway that includes the superior colliculus to the inferior olive [22,23]. As the primary saccade concludes and the target is not on the fovea, but say at 3°, the 3° region of the colliculus is unexpectedly activated [24], resulting in a secondary (corrective) saccade. However, the unexpected collicular activation is likely to also engage the inferior olive, which in turn modulates the probability of complex spikes in the cerebellum. Therefore, the secondary saccade is a motor error that induces learning. Indeed, here we found robust evidence for this trial-to-trial learning: following a secondary saccade, the velocity of the next primary saccade was adjusted in the direction of the motor error.

In our experiment, we were able to alter the properties of the secondary saccade, increasing its vigor via +RPE events and decreasing it via -RPE events. This change in vigor is likely to have been due to changes in collicular activity, resulting in earlier and slightly stronger collicular burst in +RPE trials, and later and somewhat weaker collicular burst in -RPE trials. It is possible that this RPE dependent modulation of colliculus then led to slight modification of the olive’s response, resulting in changes to the probability of complex spikes that were produced in the cerebellum. Although we did not see a significant effect of RPE on learning from movement errors, the effects that we observed were in the expected direction, with +RPE inducing a slightly greater learning than -RPE. it is possible that further research regarding use of RPE to indirectly modulate probability of complex spikes may provide an effective method for increasing motor learning rates.

### Limitations

Our interpretations regarding RPE events rely on the assumption that the opportunity to gaze at an image serves as proxy for reward acquisition. This assumption is based on the observation that for humans, gazing at images follows many of the behavioral characteristics associated with acquisition of primary rewards (i.e., food). For example, people make saccades that are faster toward images that they prefer [3], they gaze for a longer period of time at those images [10], and are willing to pay a greater effort cost in order to have the opportunity to view their preferred images [10]. Furthermore, viewing of images activates the reward system of the brain [11]. However, despite these observations, the question of whether gazing at an image engages the dopamine system remains to be explored.

In summary, whereas earlier work had demonstrated that movements are more vigorous toward more rewarding stimuli, here we found that the RPE event that takes place in the moments before onset of a movement, and not reward in itself, is necessary for modulation of movement vigor.

## Methods

A total of n=55 healthy subjects (18-41 years of age, 23±7 mean±SD), 34 females) participated in this study. The procedures were approved by the Johns Hopkins School of Medicine Institutional Review Board. All subjects signed a written consent form.

### Data collection procedure

We asked whether an RPE event systematically altered the vigor of the ensuing movement. To answer this question, we presented subjects with a visual image, and then waited until they made a saccade to it. During this primary saccade, we moved the image to a new location, and on occasion, changed its contents. The change in image location resulted in a secondary saccade. We hypothesized that the change in image content produced an RPE event. Our hypothesis was that the vigor of the secondary saccade would be affected by the magnitude of the RPE event. If the new contents were better than expected, resulting in a +RPE event, the vigor of the following saccade should be increased.

Subjects sat in front of an LED monitor (27-inch, 2560×1440 pixels, light gray background, refresh rate 144 Hz) placed at a distance of 35 cm while we measured their eye position at 1000 Hz (Eyelink 1000). Each trial began with presentation of a fixation spot (a green dot, 0.3×0.3 deg) that was randomly drawn near the center of the screen (the fixation spot was placed in a virtual box at −3 to +1 deg along the horizontal axis, and −1.5 to +1.5 deg along the vertical axis, where 0,0 refers to center of the screen). After a random fixation interval of 250-750ms (uniform distribution), the fixation spot was erased and a primary image was placed at 9 deg to the right along the horizontal axis. The size of this image was constant for each subject, but varied between subjects: 1.5×1.5 deg for some (n=20), 3.0×3.0 deg for others (n=35). A green fixation dot always appeared at the center of every image. As all analyses relied on within-subject effects, data were combined in these two groups.

The removal of the central fixation dot and presentation of the primary image served as the go signal for the primary saccade. This saccade was detected in real-time via a speed magnitude threshold of 20 °/s, or an eye position change of 2 deg from fixation, whichever happened first. After detecting saccade onset, the primary image was erased and a new image was displayed at a distance of 3 deg from the original image. A green fixation dot also appeared at the center of the secondary image. As a result, following the completion of the primary saccade, subjects produced a secondary saccade to the secondary image. The location of the secondary image was random on each trial. The reason for this was to preclude accumulation of adaptation on the primary saccades that results from movement errors that subjects experience on every trial. For some subjects (n=33), the secondary image was randomly located at either +3 or −3 deg along the horizontal axis with respect to the primary image. For other subjects (n=22), the secondary image was randomly located at each trial at either +3 or −3 along the vertical axis with respect to the primary image. As a result, the location of the secondary image was random along the horizontal or vertical axis. The size of the secondary image was always the same as the primary image.

Following completion of the secondary saccade, subjects were provided with 250ms to view that image. At the end of this period the image was erased and a center fixation dot appeared at a random location near the center of the screen, in the bounding box defined above. Each session contained 13 blocks of trials, with 100-145 trials per block. Subjects were provided with a 30 second rest period between each block.

Images were chosen from two categories: faces and noises. The facial images were gathered from the Internet (500 total images) and were modified in a way that the center of the eyes positioned at the center of the image. The noise images were constructed by shuffling the pixels of each face image (500×500pixels). This ensured that the luminance and color content of the two categories were identical.

### Magnitude of the RPE event

In our experiment we presented a primary image (e.g., face), and then at random trials replaced it with another image (noise). We hypothesized that viewing each image was a rewarding event, and as a result a difference between the primary and secondary images would produce a reward prediction error prior to the execution of the secondary saccade. We estimated the magnitude of the RPE from probability of each image and its relative value.

An objective estimate of the value of a face image with respect to a noise image can be attained from the choices that people make when given the option of viewing these images. For the specific images that we employed here, people on average chose the face image twice as often as the noise image [10]. This suggests that the relative value of face to noise is around 2.

On a given trial, the primary image was changed with a probability of 50%. Assuming a prior probability that target of the saccade is unlikely to change, and the observed likelihood that on 50% of trials the primary image changes, we can write the predicted value of the primary image as follows:

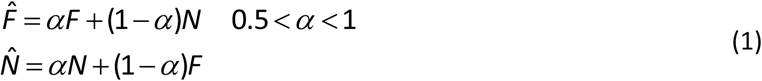

In the above expression, *F* and *N* represent the subjective value of face and noise images, and 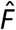 and 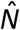 are the predicted value. Because the primary image can change, the first equation in the above expression implies that the predicted value for a primary face image is less than its subjective value (the face can become noise). The second equation implies that the predicted value for a primary noise image is greater than its subjective value (noise can become face).

Once the primary saccade concludes, the subject is presented with a secondary image. This is the image that they will actually have the opportunity to gaze at. We define RPE as the value of the second image (reward we will receive) minus its predicted value (reward we had predicted). For example, if *A* is the primary image, and *B* is the secondary image, then *Â* is the reward predicted, but *B* is the reward that will be received. That is, *RPE* [*A,B*] = *B* – *Â*.

There are four possible pairs of primary and secondary images. For each pair, we can compute the RPE:

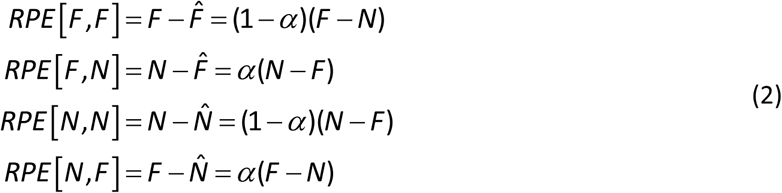

If we assume that the subjective value of a face image is roughly twice that of noise, *F ≈* 2*N*, we have:

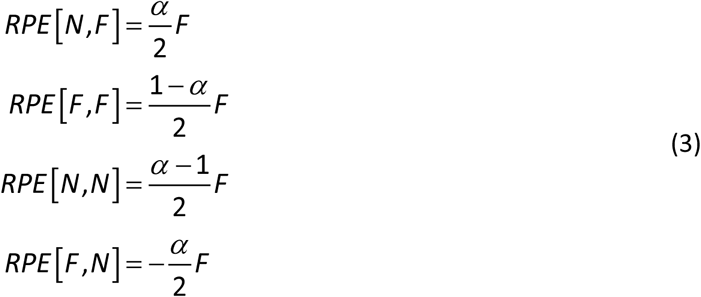

The above expressions imply the following: a noise-face trial is a very positive RPE event (first equation in the above expression), a face-face trial is a mildly positive RPE event (second equation), a noise-noise trial is a mildly negative RPE event (third equation), and a face-noise trial is a highly negative RPE event (fourth equation).

If vigor is modulated by RPE, our analysis predicts that the secondary saccades should exhibit their highest vigor in NF trials, and lowest vigor in FN trials. In comparison, FF trials should show smaller vigor as compared to NF trials, despite the fact that in both trials the secondary saccade is toward a face. Finally, NN trials should show a greater vigor than FN trials, despite the fact that in both trials the secondary saccade is toward noise.

### Data analysis

Eye position data were filtered with a second-order Butterworth low-pass filter with cutoff frequency of 100 Hz. Eye velocity data in offline analysis were calculated as the first derivative of the filtered position data. Saccades were identified with a speed magnitude threshold of 20°/sec, and minimum hold time of 10 ms at saccade end (i.e. velocity magnitude could not exceed the cutoff for a minimum 10 ms after endpoint). We measured reaction-time of the secondary saccade via the time period between offset of the primary saccade and onset of the secondary saccade. Secondary saccades onset and offset were detected identically to the primary saccades, using 20°/sec threshold on velocity magnitude. The saccade duration was considered as the time between saccade onset and offset.

Statistical analyses were performed using SPSS and general linear models, with stimulus value (e.g., face or noise) serving as the within-subject factor. We reported results of tests of within-subject effects under the assumption of sphericity. We tested this assumption via Mauchly’s test, which was confirmed in every case reported. We also performed two-sided t-tests on the between-subject effect of learning from movement errors.

## Acknowledgements

The work was supported by grants from the NIH (5-R01-NS078311), the Office of Naval Research (N00014-15-1-2312), and the National Science Foundation (CNS-1714623).

